# Holistic face recognition is an emergent phenomenon of spatial integration in face-selective regions

**DOI:** 10.1101/2020.10.13.338491

**Authors:** Sonia Poltoratski, Kendrick Kay, Dawn Finzi, Kalanit Grill-Spector

**Author notes:** corresponding author. Building 420, 450 Jane Stanford Way, Stanford, CA 94305.

## Abstract

Spatial processing by receptive fields is a core property of the visual system. However, it is unknown how spatial coding in high-level regions contributes to recognition behavior. As face inversion is thought to disrupt typical ‘holistic’ processing of information in faces, we mapped population receptive fields (pRFs) with upright and inverted faces in the human visual system. In face-selective regions, but not primary visual cortex, pRFs and overall visual field coverage were smaller and shifted downward in response to face inversion. From these measurements, we successfully predicted the relative behavioral detriment of face inversion at different positions in the visual field. This correspondence between neural measurements and behavior demonstrates how spatial integration in face-selective regions enables holistic processing. These results not only show that spatial processing in high-level visual regions is dynamically used towards recognition, but also suggest a powerful approach for bridging neural computations by receptive fields to behavior.

---

Visual recognition is critical to human behavior, informing not only what we see, but also how we navigate, socialize, and learn. In humans and non-human primates, visual recognition is carried out by a series of hierarchical, interconnected cortical regions spanning from the occipital pole to the ventral temporal cortex (VTC^1–3^). At the early cortical stages of this ventral visual stream, neurons respond to spatially local regions in the visual field and are typically tuned to simple stimulus properties like orientation or spatial frequency^4^. Later regions in the hierarchy show selectivity for complex stimuli like faces or places, and their responses correspond to perceptual experience^5,6^. These responses are thought to be abstracted from and largely invariant to low-level image properties like stimulus size or position in the visual field^7,8^. However, modern research suggests that neural responses in VTC are sensitive to both stimulus size and position^9–14^ providing a key empirical challenge to classical theories. Yet, it remains unknown if and how spatial processing in high-level visual regions is used toward recognition behavior.

The basic computational unit of sensory systems is the *receptive field* (RF), the region in space to which a neuron responds^4^. Throughout the visual system, neurons with similar RFs in retinotopic space are clustered in cortex, allowing researchers to use fMRI to measure the *population receptive field* (pRF), or the portion of the visual field that is processed by the population of neurons in a voxel^15^. Recently, our group and others have used pRF modeling to quantify spatial processing in face-selective VTC regions, finding that pRFs in face-selective regions are progressively larger than in early visual areas and are densely centered around the center of gaze (the fovea), resulting in foveally-biased coverage of the visual field^16–18^. These response properties may enable neurons in face-selective regions to integrate information across facial features for effective face recognition. Indeed, pRFs and the resulting visual field coverage in face-selective regions become larger and more foveal across childhood development, alongside with changes in fixation patterns and improved recognition of faces^17^. These findings lead to the novel hypothesis that spatial processing by pRFs in high-level visual regions may support recognition behavior.

To test this hypothesis, we examined the core human visual face network^19^ (**Figure 1a**), here defined as four anatomically distinct functional regions: one in the inferior occipital gyrus (IOG-faces/OFA), one on the posterior fusiform gyrus (pFus-faces/FFA1), one straddling the mid-fusiform sulcus (mFus-faces/FFA2), and one on the posterior end of the superior temporal sulcus (pSTS-faces). This network serves as an ideal model system for deriving links between neural computations and behavior, as it is reliably localized in each human participant, causally involved in face recognition behavior^20,21^, and stereotypically organized in relation to known anatomical landmarks and cytoarchitectonic regions^22^. Importantly, face recognition behavior has been richly characterized across decades of psychology research^23,24^, allowing us to link neural responses to robust and replicable behavioral markers of face recognition.

**Figure 1.**
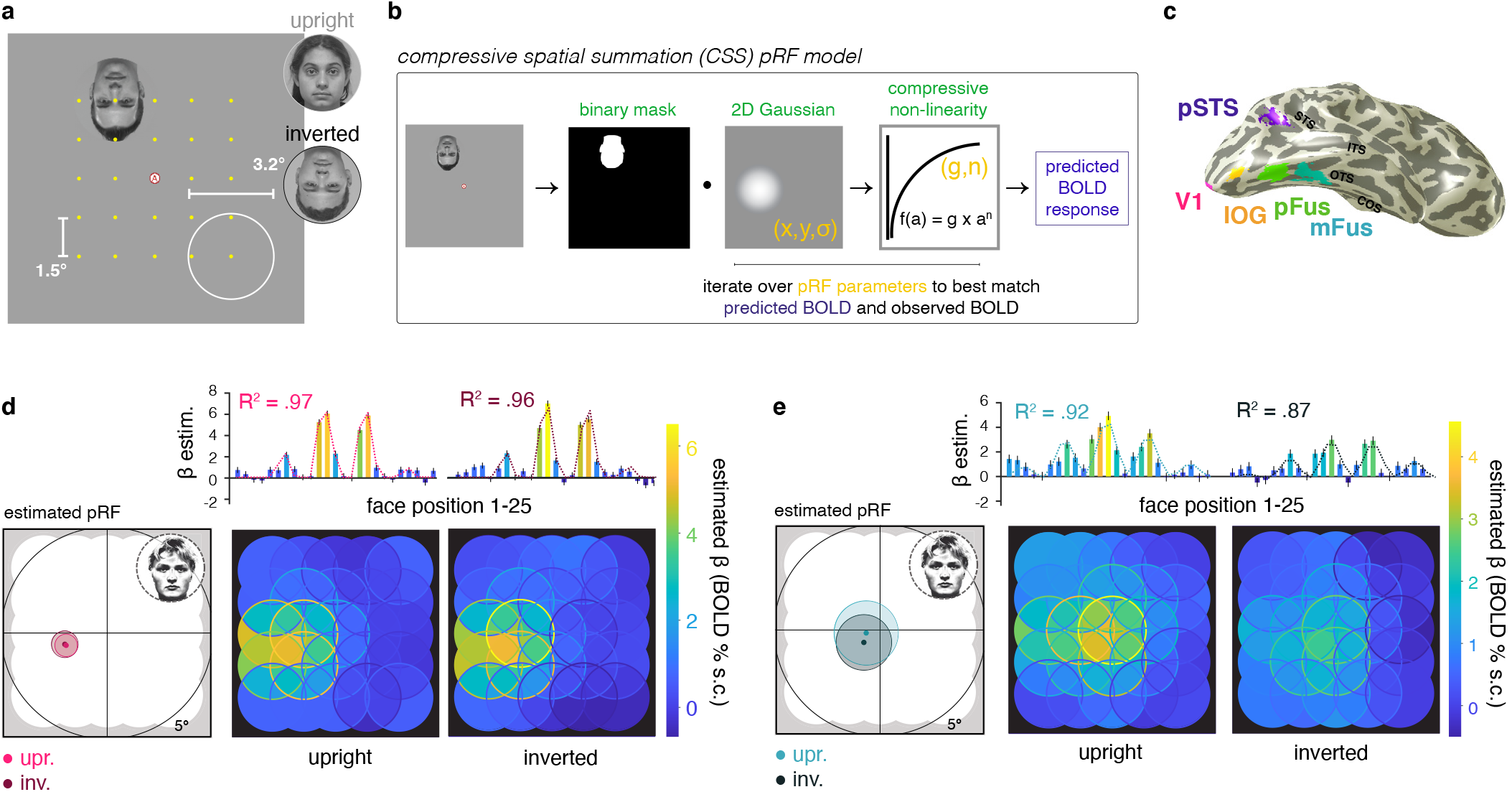
Experimental design and pRF model fitting procedure. **(a)** Schematic illustrating the 5 × 5 grid of positions (1.5° center-to-center) at which faces (3.2° diameter) were presented, either upright or inverted as shown. **(b)** Model fitting procedure of the compressive spatial summation (CSS^29^) pRF model, which represents the position of each face in the visual field as a binary mask. Estimated parameters (*yellow text*): center position X and Y, spread σ, exponent (n), and gain (g). While a single binary mask is shown here, an averaged silhouette mask of the faces that each participant saw at each position was used for the model fitting. **(c)** V1 and face-selective regions of interest on the cortical surface of a representative participant. *Acronyms*: V1: primary visual cortex; IOG: inferior occipital gyrus face-selective region, also referred to as the occipital face area (OFA); pFus: posterior fusiform face-selective region, also referred to as fusiform face area 1 (FFA-1); mFus: mid fusiform face-selective region, also referred to as fusiform face area 2 (FFA-2); pSTS: posterior superior temporal sulcus face-selective region. **(d, e)** Example V1 and mFus-faces voxel data and model fits. *Top* (*bar plots*): BOLD responses (β estimates in % signal change) for each of the 25 mapping positions for this voxel, plotted first from left to right, and then top to bottom. *Error bars*: ± SEM across runs; *Dotted lines*: pRF model fit, which was estimated separately for upright and inverted faces. *Left*: estimated pRF in response to upright (*light colors*) and inverted (*dark colors*) faces. Circles contours indicate estimated pRF size, radius = σ / √n. *Right* (*overlapping-disk*) *panels*: Responses at each position at which the mapping stimuli appeared, colored by response amplitude. Overlapping stimuli responses are averaged within each disc, while the absolute responses at each position are indicated by the disc outline.

Broadly, we hypothesized that computations by pRFs in face-selective regions provide the basis for spatial integration of information toward face recognition behavior. This led us to predict that face inversion, which is thought to hinder typical spatial processing of faces^23,25–27^, would alter pRF estimates in face-selective regions, but not in early visual regions where spatial integration is not sensitive to face content. To test our hypotheses, we mapped pRFs across the ventral visual pathway with both upright and inverted faces (**Figure 1b**). We predicted that if large and foveal pRFs in face-selective regions are adaptive for typical recognition behavior, both individual pRFs and the visual field coverage afforded by a cortical region may be shifted in position and/or in size with face inversion. Alternatively, if spatial coding in face-selective regions is a concomitant neural property that does not actively play a role in face recognition, we would expect equivalent estimates when mapping with upright or inverted faces in both face-selective regions and early visual cortex.

Finally, we hypothesized that if pRFs reflect neural mechanisms underlying functional face recognition, they may link directly to the behavioral face inversion effect (FIE^23^. First described over 50 years ago, the FIE is a robust and specific behavioral deficit in recognizing faces when they are presented upside down. Critically, the FIE has been associated with a failure of typical spatial pooling of information across multiple features of a face^24,28^, prompting us to examine the linkage between the visual field coverage of face-selective regions and the behavioral FIE. We reasoned that if pRF coverage in face-selective regions is reduced or shifted in response to face inversion, then neural processing of inverted faces at the center of the visual field would be suboptimal and behavioral recognition would be impaired. Accordingly, the behavioral FIE may be mitigated if faces are placed in the optimal location for inverted faces as determined from the visual field coverage they elicit. As we found that pRFs and the visual field coverage in face-selective regions where smaller and shifted in response to inverted faces, we tested these predictions in a separate behavioral experiment outside the scanner by using our participants’ neural measurements to predict the magnitude of the FIE across visual field positions.

## Results

To map population receptive fields in response to upright and inverted faces, twelve participants took part in an fMRI experiment. Faces were presented in randomized order at 25 locations spanning the central ~9.2° of the visual field while participants maintained fixation on a central letter stimulus (**Figure 1a**; Online Methods); fixation is critical for these measurements to ensure that the retinotopic position of the stimulus can be reliably coded in the model. During pRF mapping, participants performed a challenging rapid visual stimulus presentation (RSVP) letter task at fixation. While they were aware that faces would appear on the display, participants were instructed that faces were not task-relevant during the fMRI session. We saw no differences in behavioral performance on the letter task between trials in which upright (90.2 ± 1.8% correct) or inverted (89.4 ± 2.5%) faces were presented (*t*(11) = 0.531, p = 0.606). Therefore, we infer that any observed differences in pRFs measured in response to upright and inverted faces are stimulus-driven by the face inversion, rather than by task performance or attention.

We implemented the compressive spatial summation (CSS) pRF model^16,29^, which models each voxel’s pRF as a circular Gaussian with a compressive nonlinearity (n; **Figure 1b**). Each model fit yields five parameters that describe the gain (g), position (x,y), and size (σ/ön) of the pRF of each voxel, as well as a measure of the goodness-of-fit of the model (R^2^). pRFs were estimated independently for upright and inverted faces in each voxel in the primary visual cortex (V1) and four face-selective cortical regions (**Figure 1c**); data from V1 is presented as an early visual control. As illustrated for example voxels from V1 (**Figure 1d**) and mFus-faces (**Figure 1e**), the position of the face in the visual field robustly modulates the response of these regions. Further, the CSS pRF model accurately quantifies these voxels’ responses, as it explains the majority of their respective variances (**Figure 1d,e-R^2^**). Notably, the independent model fits yield similar pRF estimates across upright and inverted faces in V1, corresponding to predictions that spatial responses in early visual cortex are largely agnostic to this manipulation of stimulus content (**Fig 1d-left panel**). However, for the example mFus-faces voxel, face inversion yields responses that are lower in magnitude and shifted in the visual field, which consequently produce differential pRF estimates across the two mapping conditions (**Fig 1e-left panel**).

### Mapping with inverted faces modulates pRF properties in face-selective regions but not V1

We first quantified and compared average pRF properties for upright and inverted faces in face-selective regions. We found no significant differences in the horizontal position of pRF centers across upright and inverted mapping conditions in face-selective regions (**Figure 2a, Table S1**). However, face inversion yielded pRF centers that were consistently shifted downwards (**Figure 2b**). We quantified changes in estimated pRF parameters using a 3-way repeated measures analysis of variance (ANOVA) with factors of ROI (IOG-/pFus-/mFus-/pSTS-faces), hemisphere (right/left), and mapping condition (upright/inverted). This revealed a significant effect of inversion on pRF center Y position (F(1,10) = 8.82, p = 0.014; no other significant main effects or interactions, **Table S1**). Post-hoc t-tests revealed that downward shifts in the average pRF position were significant in all ventral face regions (IOG-, pFus-, mFus-faces, **Figure 2b**). Additionally, pRF gain was significantly lower for inverted than upright faces in some of the face ROIs (**Figure 2c**; significant ROI by inversion interaction F(3,30) = 4.66, p = 0.009, 3-way repeated-measures-ANOVA; no other significant effects, **Table S1**). Post-hoc t-tests showed that pRF gains were lower for inverted than upright faces in IOG- and mFus-faces (**Figure 2c**). This finding is consistent with prior reports of lower mean responses for inverted than upright faces in ventral face regions^25,26,30^.

**Figure 2.**
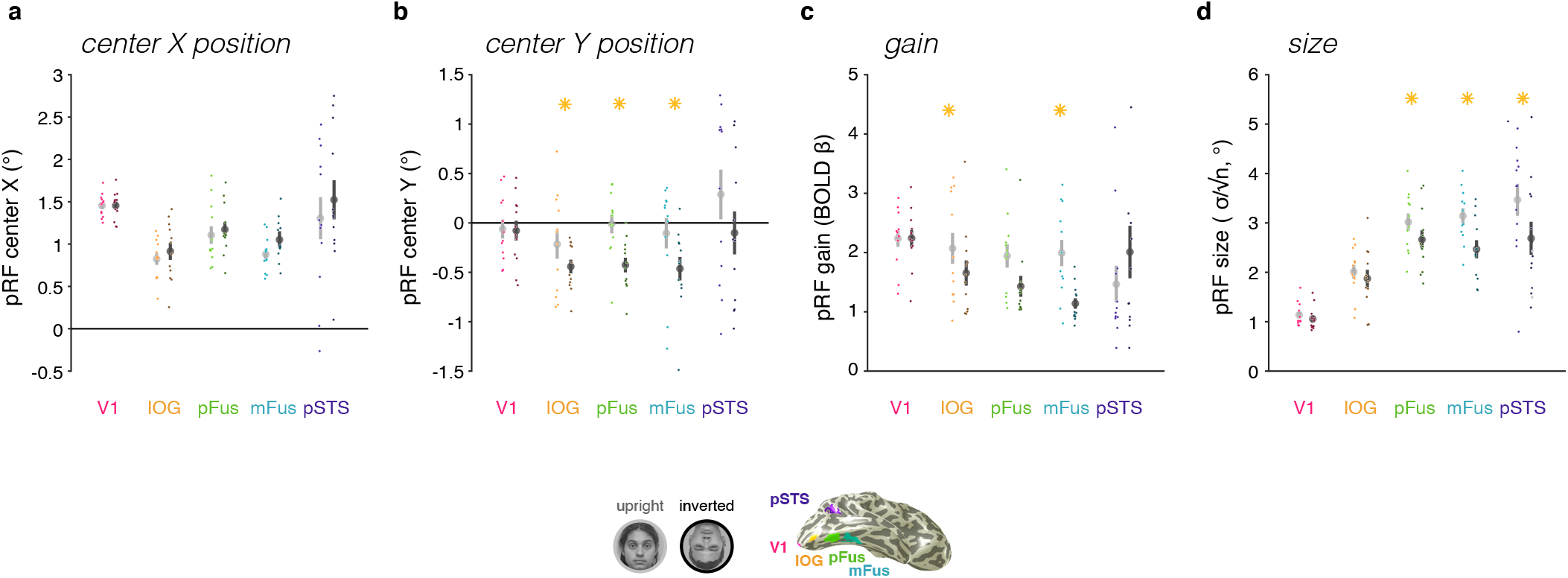
Face inversion impacts pRF estimates in face-selective cortical areas but not primary visual cortex. Mean pRF properties when mapped with upright faces (*light grey*) and inverted faces (*dark grey*) in bilateral regions of interest across 12 participants. *Error bars*: ± SEM across participants. *Dots*: individual participant means. *Asterisks*: statistically significant differences between upright and inverted face mapping conditions (two-tailed paired t-test across participants, p < .05). **(a)** pRF center X position; right hemisphere data are mirror flipped to left. **(b)** pRF center Y position. **(c)** pRF gain. **(d)** pRF size (σ / √n).

Interestingly, there were also significant differences in estimated pRF size across upright and inverted mapping conditions, which varied by face-selective region (ROI by inversion interaction F(3,30) = 7.53, p = 0.0001, 3-way repeated-measures-ANOVA, **Table S1**). Specifically, pRF size was significantly smaller in fusiform face-selective regions (pFus- and mFus-faces) and pSTS-faces in response to face inversion (*t*’s(11) > 2.98, *p*’s < 0.013; **Figure 2d**), but not in IOG-faces.

Overall, face inversion had the most pronounced effect on pRF properties in mFus-faces, the end stage of the ventral face-processing hierarchy^18,19^. In mFus-faces, face inversion yielded a significant downward shift of pRFs, smaller pRF size, and smaller gain. As we did not find any significant difference across hemispheres of the effect of inversion on pRF responses (**Table S1**), we combine data across hemispheres unless noted. In contrast to the profound effect face inversion had on pRF properties in face-selective regions, we saw no significant differences between mapping conditions in any estimated pRF property in V1 (**Figure 2-V1**). This suggests that spatial integration is altered in response to face inversion specifically in higher-level face-selective regions, but not in early visual cortex.

### Signal-strength differences are insufficient to explain the observed effects

As BOLD amplitudes were generally reduced in face-selective regions in response to inverted faces, we observed lower estimated pRF gain (**Figure 2c**). Additionally, model goodness-of-fit in face-selective regions was consistently lower in the inverted than in the upright mapping condition (**Figure S1**). We asked if these reductions in signal strength might explain the observed differences in pRF position and size. To test this possibility, we simulated the effect of changing the magnitude of responses and level of noise in mFus-faces on model estimates of pRF center Y position and size. Results of the simulations suggest that differences in signal strength differences are insufficient to account for the scale of our observed effects in mFus-faces (**Figure S2b-c**).

### pRFs in face-selective regions are smaller across eccentricities in response to inverted faces

The analyses of mean pRF size are consistent with the hypothesis that face inversion yields more spatially constrained integration of information via pRFs in face-selective regions. However, as pRF size linearly increases with eccentricity throughout the visual hierarchy^15–16^, it is critical to consider the relationship between size and eccentricity when evaluating pRF size estimates across conditions. Thus, we compared the size vs. eccentricity relationship for upright and inverted faces in each ROI. We reasoned that if there are differences in spatial processing across mapping conditions,we should observe a higher intercept or a steeper slope relating size to eccentricity for pRFs mapped with upright than with inverted faces.

As evident in **Figure 3**, we observed smaller pRF sizes in response to inverted than to upright faces across eccentricities in pFus- and mFus-faces. In contrast, we saw minimal differences in pRF sizes across eccentricities between upright and inverted face mapping for either IOG-faces or V1, consistent with data in **Figure 2d**. To evaluate these results across participants, we performed an equivalent line-fitting separately for each participant (**Figure S3**) and tested for significance. Results indicate the slope relating pRF size and eccentricity is not affected by face inversion (F(1) = 0.04, p = 0.852, 2-way repeated measures ANOVA on slope with factors of ROI (IOG-/pFus-/mFus-/pSTS-faces) and condition (upright/inverted; **Table S2**). However, pRFs were smaller across eccentricities (significant main effect of intercept (F(1) = 9.08, p = 0.012, 2-way repeated measures ANOVA, **Table S2**). Post-hoc t-tests revealed that these effects were significant in both pFus- and mFus-faces (pFus: *t*(11) = 4.185, p = 0.002; mFus: *t*(11) = 2.371, p = 0.037) but not in IOG-faces, pSTS-faces, or V1.

**Figure 3.**
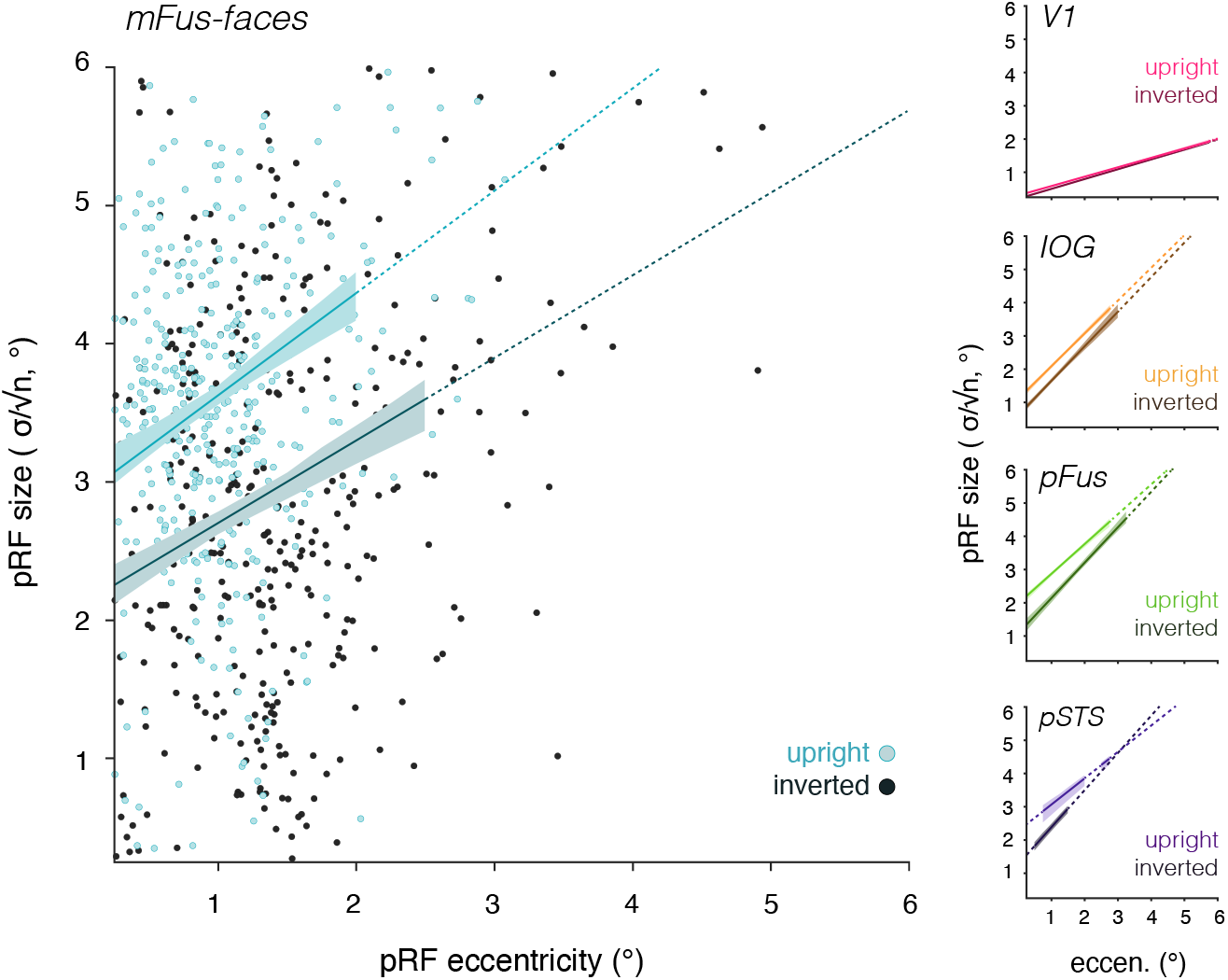
Face inversion yields smaller pRFs in face-selective regions across eccentricities. The relationship between size and eccentricity in each ROI, for voxels mapped with upright faces (light colors) and inverted faces (dark colors). This relationship is summarized by the line of best-fit across all voxels pooled over participants with model goodness of fit R^2^ > .5. *Shaded error*: a bootstrapped 68% confidence interval at bins (0.25° wide) for which there were more than 20 voxels across participants. *Dashed lines*: bins for which there were fewer than 20 voxels across participants. *Dots*: a random sample of 300 mFus-faces voxels across participants. See also **Figure S3**.

### pRFs in face-selective regions are shifted downward in response to inverted faces

To further explicate the effect of face inversion on pRF position in face-selective regions, we calculated the distributions of pRF positions in each participant and ROI when mapped with upright and inverted faces (**Figure 4**). In IOG-faces, the average distribution of pRF center Y positions mapped with upright (**top panel**) and inverted (**bottom panel**) faces were largely equivalent (**Figure 4a**). In contrast, face inversion yielded a robust downward shift in pFus-, mFus-, and pSTS-faces. In fusiform face-selective regions, inversion shifted the overall distributions’ centers from the horizontal meridian to the lower visual field. These shifts are visible when plotting the location of each voxel’s upright vs. inverted pRF position as shown in **Figure 4b** for mFus-faces. While we also see some variability in the estimated pRF center X position across upright and inverted mapping, these shifts have no consistent direction, even when considering hemispheres independently (**Table S1, Figure 2a**). The downward shift in pRF center position in response to inverted faces altered the overall distribution of pRFs to be further from the fovea. Notably, increased eccentricities would have predicted larger pRFs mapped with inverted than with upright faces, which is the opposite of our empirical findings (**Figure 3**).

**Figure 4.**
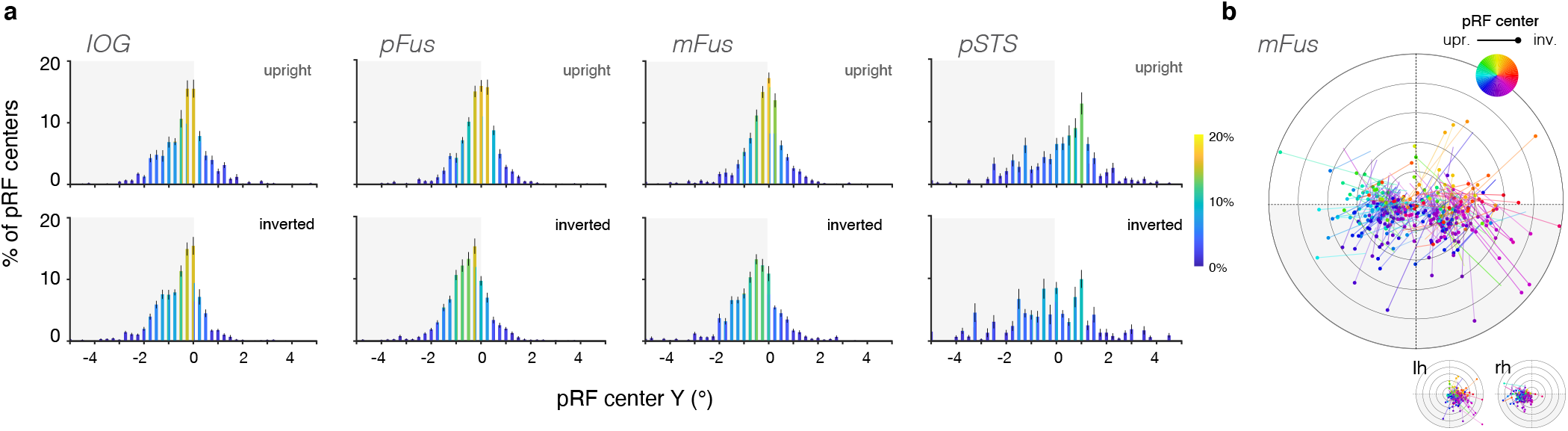
In face-selective regions, inverted-mapped pRFs differ in vertical position Y. **(a)** Across participants, the average distribution of pRF position Y when mapped with upright (top row) and inverted (bottom row) faces. *Error bars*: ± SEM across participants in each 0.25° bin. **(b)** Position shifts for a random subsample of 300 mFus-faces voxels across participants (for which the model variance explained is higher than 50% of the variance); voxel pRF center position estimates are consistently shifted downward.

### pRF changes result in differences in the coverage of the visual field in face-selective regions

Given our findings that pRFs in face-selective regions are modulated by face inversion, we next asked how these changes may impact the overall coverage of the visual field afforded by these regions. This visual field coverage reflects the distributed spatial responsivity of the entire population of neurons a cortical region, and thus may be more closely linked to the behaviors that the population subserves. Thus, we constructed and quantified density coverage maps in each ROI for each participant and then averaged across participants (see Online Methods, **Figure 5**).

**Figure 5.**
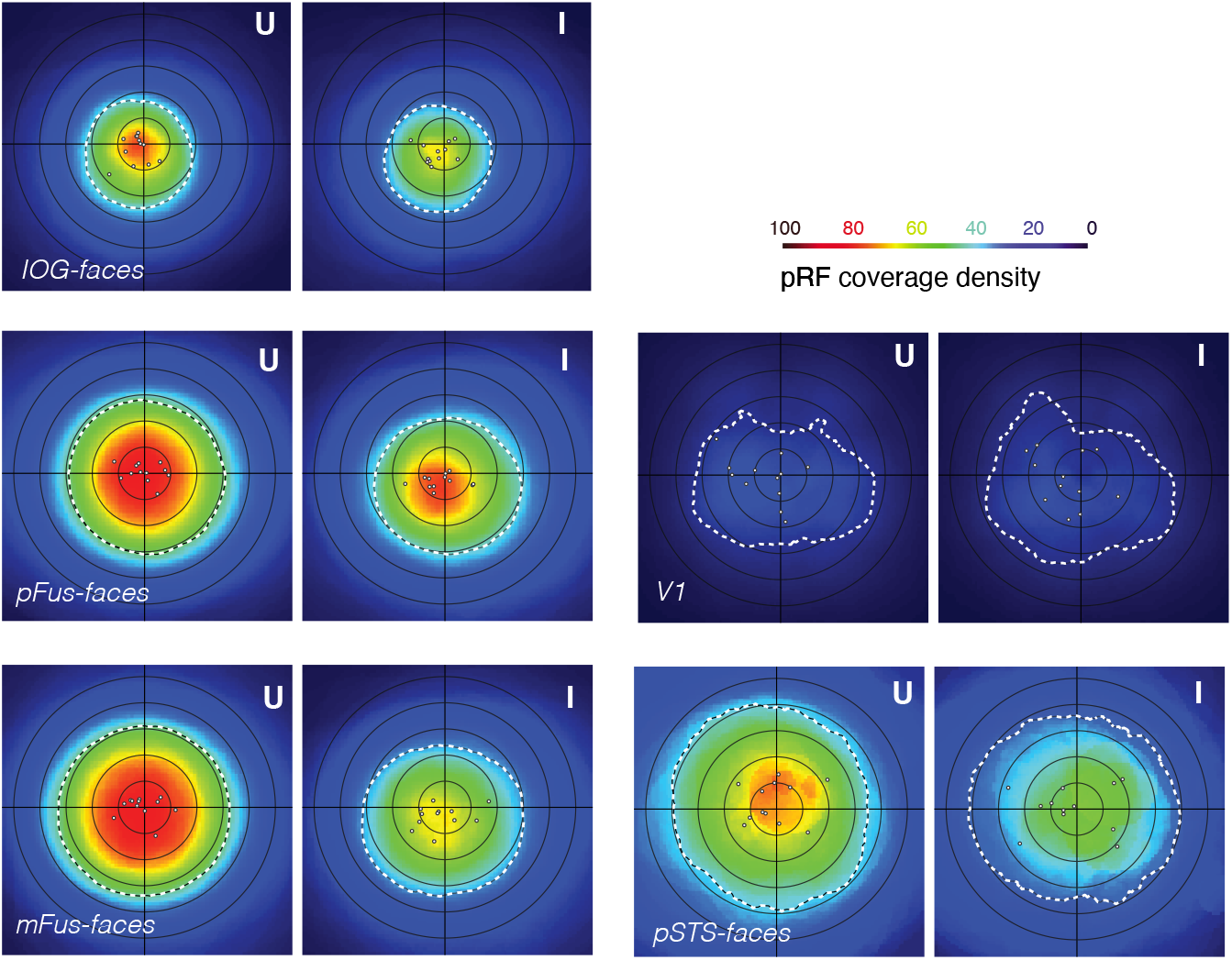
Face inversion leads to differences in overall visual field coverage in face-selective regions. **(a)** Average pRF density plots for each region across participants (N = 12) when mapped with upright (left) and inverted (right) faces. Density plots are bootstrapped for each participant using 1000 draws of 80% of the voxels and then averaged across participants (see Online Methods). The resulting plot color indicates the average percentage of voxels whose pRFs cover each point in retinotopic space, where coverage is defined as within 2σ /√n ° from the pRF center. *Dashed white contours*: full width half maximum (FWHM) of the coverage density, which we use as a summary of the overall coverage field of view of the region. *Dots*: coverage center of mass of each individual participant.

When pRFs are mapped with upright faces, ventral face-selective regions show a prominent foveal bias in coverage (**Figure 5-left-U**). This foveal bias is markedly reduced in response to inverted faces (**Figure 5-left-I**), and is coupled with a significant reduction in the coverage area and downwards shift of the coverage (**Table S3, Figure S4**). These effects appear largely consistent across individual hemispheres (**Figure S5**). As in voxelwise analyses, mFus-faces shows the strongest differentiation between upright- and inverted-mapped coverage (**Figure 5**). Coverage for the latter spans a smaller portion of the visual field (significant change in full-width-half-max area: *t*(11) = 2.79, p = 0.017) and is shifted downward relative to the former (significant change in center-of-mass Y: *t*(11) = 3.338, p = 0.007, but not center-of-mass X: *t*(11) = −0.569, p = 0.58, **Figure S4**). A similar downward shift is evident in pFus-faces (significant change in center-of-mass Y: *t*(11) = 3.123, p = 0.01). In contrast, IOG-faces, pSTS-faces and V1 showed minimal differences in coverage between the mapping conditions.

Given prior reports of stimulus-dependence of visual field coverage in nearby word-selective regions^31^, we sought to examine whether differences in coverage in face-selective regions were specific to inversion. We evaluated this by measuring coverage in our participants’ face-selective regions to different stimuli: brightly colored cartoon images in a separate pRF mapping experiment (**Figure S5a,** ^18^). Coverage in face-selective regions mapped with these highly varied stimuli was similar to the upright faces condition; pRF density was highest around the fovea, and consequently the coverage was centered on the fovea (**Figure S5b-c**). This suggests that reduction in area and the downward shift of coverage observed for inverted faces is specific to face inversion, and does not occur for any suboptimal mapping stimulus.

In all, we see profound differences in the way that face-selective regions sample visual space in response to face inversion, both at the level of individual voxels, and by the collection of voxels constituting a region. Importantly, pRF differences are observed even in response to passively-viewed mapping faces, as participants’ attention was diverted to an unrelated RSVP task on letters at the central fixation. This is particularly interesting as it suggests that the reported effects of inversion on spatial processing in face-selective regions is not a simple reflection of the participants’ task or focus of attention, but instead reflect an underlying sensory property that may ultimately constrain behavior. To evaluate this idea, we next used our fMRI measurements to derive and test a specific prediction for participants’ performance in recognizing upright and inverted faces outside of the scanner.

### A novel test of the relationship between pRF coverage and face recognition behavior

During typical face viewing, pRF coverage in face-selective areas seems optimized to capture information from the face: large and foveal pRFs extend into the contralateral visual field, allowing for the processing of many features of a centrally-viewed face by the pRFs of a face-selective region^16^. However, as schematized in **Figure 6a**, our data suggest that this is not the case when viewing inverted faces. If participants fixate at the center of an inverted face in a standard recognition task, our data predict that the visual field coverage by pRFs of face-selective regions involved in recognition will suboptimally cover the inverted facial features (**Figure 6a**). This leads to an intriguing hypothesis about the functional role of spatial integration in face-selective regions: if pRF coverage corresponds to windows of spatial integration toward face recognition, then increasing the overlap of this coverage with inverted facial features should yield better recognition performance of inverted faces, consequently reducing the magnitude of the behavioral face inversion effect (FIE) in that location in the visual field.

**Figure 6.**
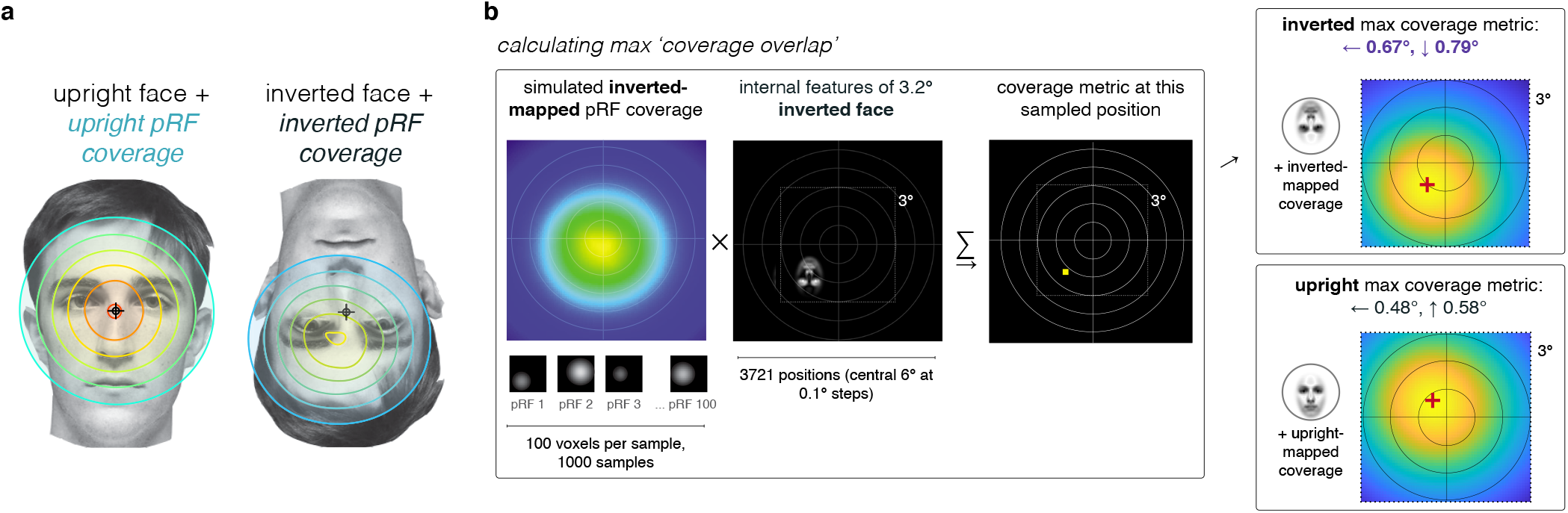
Using visual field coverage of face-selective regions to predict the location of improved performance for inverted faces. **(a)** Schematized depiction of upright- and inverted-mapped mFus-faces pRF coverage when one views example faces centered on fixation. This predicts that viewing inverted faces yields suboptimal coverage of the location of the face features by pRFs in face-selective regions, which may contribute to the reported behavioral detriments in recognizing inverted faces (e.g. the face inversion effect, or FIE). **(b)** To evaluate the hypothesis schematized in (a), we used a simulation procedure (see Online Methods) to determine the location in the visual field that would produce maximal pRF overlap with the internal features of 3.2° faces. To do so, we iteratively sampled a subset of mFus-faces voxels to generate bootstrapped coverage in response to inverted or upright faces; we then took the dot product of this bootstrapped coverage and a face centered at various retinotopic positions (central 3° × 3°, sampled at 0.1° steps). The retinotopic location that yielded the maximal overlap (dot product) between the inverted face and inverted-mapped pRF coverage, as well as the upright face and upright-mapped coverage, is shown in the right panels.

To evaluate this hypothesis, we first used a simulation to determine the location at which we expected the behavioral FIE to be attenuated (**Figure 6b**, Online Methods). As a large body of behavioral research indicates that internal facial features are key for recognition^23,25–27^, we reasoned that performance for inverted faces would be best at the retinotopic location at which its internal features maximally overlap with the inverted-mapped pRF coverage in face-selective regions. To determine this location, we simulated the overlap between the internal features of inverted and upright faces and the corresponding pRF coverage from mFus-faces, where the effects of face inversion were strongest. As in the fMRI study, we used 3.2° face images in the simulation, and determined that the optimal location to place these inverted faces was centered 0.67° leftward and 0.79° downward from fixation. Notably, this location is different than the optimal location for upright faces (**Figure 6b-bottom right**). While in both cases, predicted performance is best in the left visual field^32,33^, only performance for inverted faces is predicted to be improved in the lower visual field (see also **Figure S7**).

We tested these predictions by evaluating face recognition performance of each of our participants in a behavioral experiment conducted outside of the scanner. The behavioral study consisted of a challenging recognition memory task on upright and inverted faces (50% each, **Figure 7a**). Fixation was held at a small central bullseye, and strictly monitored (Online Methods). On each trial participants saw a sequence of 3 faces (all upright or all inverted), each shown for 400ms (**Figure 7a**). After an 800ms interval, a target face was shown, and participants were asked to indicate whether the target was present in the preceding triad (50% probability). Critically, to test our hypothesis, recognition performance on upright and inverted faces was evaluated at three retinotopic locations: the *lower left*, corresponding to maximal pRF overlap for inverted faces, *central* fixation, and an *upper right* location that was equivalent in distance from fixation (**Figure 7b**). This third condition serves as a control; if retinal acuity alone determines performance, recognition should worsen symmetrically with increasing distance from the fovea for both upright and inverted faces. Notably, these position shifts are relatively small: 3.2° faces are shifted 1.03° off-center in the lower-left and upper-right conditions. Individual trials on which participants broke fixation were excluded from the analysis, and the data of three participants who were unable to maintain consistent fixation were removed (see Online Methods).

**Figure 7.**
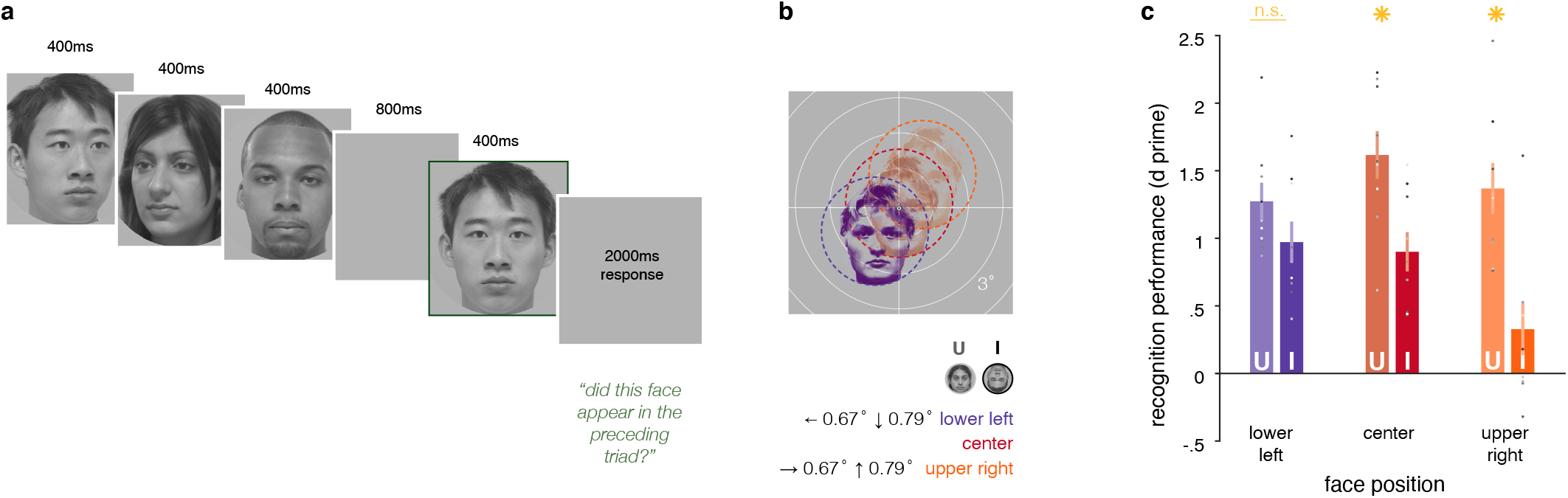
The behavioral FIE is reduced at the retinotopic position that maximally overlaps mFus-faces pRF coverage. **(a)** Illustration of the trial structure and timing of the recognition task outside the scanner, in which participants viewed a triad of rapidly-presented faces and indicated whether the subsequent target face had appeared on the trial. Within a trial, faces could be upright (as illustrated), or inverted, and appeared at the three locations described in (b). Subjects fixated on a central bullseye target (0.3° diameter), and their fixation was monitored with an eyetracker. Only trials on which participants maintained fixation were included in analyses. **(b)** Illustration of the three positions at which faces (3.2° diameter) appeared on the screen, as determined by mFus-faces pRF coverage from the fMRI experiment (**Figure 6b**). **(c)** Results of the behavioral experiment. When participants (N=9) viewed upright faces (light color bars), performance was predictably highest at the center (red), and fell when the faces were presented in the lower left (purple) or upper right (orange) positions. However, recognition performance for inverted faces (dark color bars) was highest at the lower-left location. In fact, across participants, the behavioral FIE (upright minus inverted performance) was significantly reduced in the lower-left position, while remaining strong at the center and upper-right positions. *Dots*: individual participant performance; *Asterisks*: significant differences between upright and inverted face conditions at p < .05. *N.S*.: not significant.

We found striking differences in recognition performance at the three probe locations. Recognition for upright faces was best when faces were presented at the center of the screen and declined similarly when placed at the lower left or upper right positions (**Figure 7c-U**). This is consistent with our predictions, as both the upper-right and lower-left positions were nearly equidistant from the simulated optimal position for upright faces (**Figure 6b;** the lower left is 0.22° farther). Recognition of inverted faces, however, followed a different pattern: performance was equal if not better at the lower-left location than at fixation, but starkly worse at the equidistant upper-right location (**Figure 7c-I**). Indeed, the magnitude of the FIE, or the difference between recognition performance for inverted and upright faces, was not significantly different from zero at the lower-left location (*t*(8) = 1.68, p = 0.13, 2-sided paired t-test), while remaining significant at the other locations (center: *t*(8) = 3.43, p = 0.01, upper right: *t*(8) = 5.06, p < 0.01). Thus, the behavioral FIE is attenuated by placing faces in the location where inverted-mapped coverage of the visual field in ventral face-selective regions covers facial features optimally, linking the effect of face inversion on visual field coverage in face-selective regions to face recognition performance.

## Discussion

Our experiments demonstrate stimulus-driven changes in spatial coding of face-selective regions that correspond directly to visual recognition behavior. In the neuroimaging experiment, we demonstrated that pRFs and coverage of the visual field in the face network, but not in primary visual cortex, are altered by face inversion. While each of the face-selective regions showed some degree of sensitivity to inverted faces, effects were most pronounced in mFus-faces (FFA-2), where both individual pRFs and the coverage of the visual field were shifted downward and reduced in response to inverted faces. A similar pattern of results was found in pFus-faces (FFA-1). pSTS-faces, a region in the lateral stream of the face-selective VTC network thought to process dynamic aspects of faces^34,35^, exhibited similar effects of smaller pRF size, albeit noisier responses overall. This may be attributed to the substantially more peripheral responses in pSTS-faces^18,36^, making our mapping with still images over a 9.2° × 9.2° visual field suboptimal. However, face inversion did not significantly alter visual field coverage in IOG-faces. This is consistent with the region’s strong responses to individual face features^14,37^, such as the eyes or mouth, on which inversion has a smaller perceptual effect. Importantly, we found differences in spatial processing in response to face inversion while participants performed a task that removed their directed attention from the faces. Following prior behavioral findings^38^, this suggests that bottom-up properties of the stimulus itself, and not attention or task drive the modulation of pRF properties by face inversion.

In the behavioral experiment, we demonstrated that the magnitude of the face inversion effect (FIE) varies across the visual field in a manner that is strikingly consistent with our measurements of differential visual field coverage in the fMRI study. Specifically, we showed that the magnitude of FIE can be mitigated by placing faces in an optimal location that corresponds to the inverted-mapped visual field coverage of mFus-faces. These data suggest that the FIE is a consequence of the differential population activity in face-selective regions evoked by face inversion, which corresponds to large-scale differences in the way these regions sample the visual field. More generally, our results highlight a precise relationship between spatial processing in high-level visual regions and the recognition behavior that they subserve.

### Holistic face processing as spatial integration across multiple features

An extensive body of behavioral research has shown that faces are processed holistically—that is, face recognition behavior involves concurrent processing of the whole face, rather than its individual features^23,27,28,38–42^. However, despite decades of research into the behavioral signatures of holistic processing of faces, its underlying neural mechanisms remain opaque.

From our findings, we propose that holistic processing of faces can be understood as a consequence of spatial integration of information across face features, which emerges from the large and foveal receptive fields of ventral face-selective regions. Consequently, behavioral failures of typical holistic processing, like the FIE^23^, appear to be a result of maladaptive, stimulus-driven changes of pRF coverage in these regions. These findings provide neural support to cognitive theories that predicted a reduced perceptual field for inverted faces^28^. Importantly, our data not only elucidate the neural and computational mechanisms of face inversion, but also provide for the first time a computational framework that generates quantitative predictions of behavioral deficits of holistic processing. This provides an important foundation for future work that can quantitively examine if individual differences in holistic processing^43^ and impairments in face recognition^44,45^ are derivative of differences in pRF size, position, and coverage in high-level visual regions.

### Stimulus selectivity interacts with spatial processing by pRFs

Our data demonstrate an interaction between estimates of pRF properties in face-selective regions and the stimulus used in mapping (upright vs. inverted faces), which is specific to face inversion as we did not find equivalent deviations for phase-scrambled faces^16^ or diverse cartoons^18^ **Figure S6**).

What aspect of face inversion produces the observed changes in pRFs? One possibility is that the differential location of the eyes or internal features contributes to pRF differences with face inversion. In our stimuli, the direction and magnitude of the observed Y position shift in pRF centers appears to correspond to the differential center-of-mass of the internal features between upright and inverted faces rather than solely the eyes (**Figure S8**). However, the differential location of internal features is not sufficient to explain the observed reduction in pRF size nor the lower gain. Thus, an interesting direction for future research is building an image-computable model that predicts all measured effects of face inversion. To do so, researchers may consider how the displacement of internal features may interact with the learned spatial configuration of typically-viewed, upright faces^46^, potentially yielding suboptimal contextual modulation of RFs of individual neurons^47^.

### High-level pRF processing as a bridge from brain to behavior

While spatial sensitivity has been extensively demonstrated in VTC^9–14^, the prevailing view theorizes that spatial information in ventral temporal regions is independent from the recognition that these regions support. Our findings emphasize that high-level visual regions not only maintain spatially localized processing that can be computationally modeled by pRFs, but also that this processing dynamically supports visual recognition behavior. Doing so provides the first demonstration of the functional utility of spatial processing in the ventral ‘what’ stream.

More broadly, as receptive fields (RFs) are a basic computational unit of many domains^48–52^, we propose that quantitative measurement and modeling of pRFs in high-level regions across the brain opens compelling avenues for deriving precise links between brain and behavior – especially as activity in these regions often corresponds to perceptual experience more directly than in earlier stages of neural processing. In particular, our approach can not only quantitatively determine what factors may alter pRFs in other domains, but also leverage these measurements to generate new testable behavioral predictions.

In sum, this work demonstrates a precise link between spatial processing in face-selective regions and recognition behavior. We find that the neurally ill-defined concept of ‘holistic’ face processing is an emergent property of spatial integration by population receptive fields. These data suggest promising new directions for bridging basic neural computations by receptive fields to behavior.

## Methods

### Participants

13 participants (6 women) ages 20-31 participated in a single session of pRF mapping, including authors SP and DF. Nine participants identified as white/Caucasian, 1 as Asian, 1 as Black, and 2 as mixed-race (Hispanic/white and Asian/white). One participant’s data was excluded for excessive motion (> 4mm) during the scan. All participants provided informed, written consent per the Internal Review Board of Stanford University, and were compensated for their time. No statistical modeling was used to determine the size of our participant sample as we model each voxel individually in each participant. We used typical sample sizes for fMRI pRF studies of the visual system which use data from a range of 3-20 participants^15,16,31,53–56^. All scan participants completed the subsequent behavioral experiment.

### Scanning protocol

Participants underwent a single fMRI session in the main experiment, which consisted of 8-10 runs of pRF mapping and lasted approximately 90 minutes. Scanning was done at the 3 Tesla GE research-dedicated magnet at the Stanford Center for Cognitive and Neurobiological Imaging (CNI). Functional T2* weighted pRF mapping runs were each 282s long, and collected using a 32-channel headcoil, single-shot EPI, with a voxel resolution of 2.4mm isotropic and a TR of 2s (FOV = 192, TE = 30ms, flip angle = 77°). Twenty-eight slices were prescribed parallel to the parieto-occipital sulcus to cover each participant’s occipital and ventral temporal lobes.

We additionally acquired a high-resolution (1mm isotropic voxels) T1-weighted full-brain anatomical during either the main experimental session or in a separate retinotopy/localizer session for each participant, and used this whole-brain anatomical image to align data across the experiment, localizer, and retinotopy sessions in native participant space.

### pRF mapping

Each population receptive field mapping run lasted 282 seconds, and presented face images at 25 spatial positions to span the visual field over time (**Figure 1a**). Face images were 3.2° in diameter, and were offset 1.5° center-to-center to tile approximately 9.2° × 9.2° of the central visual field. Faces were presented in 4s trials of faces (50 trials, 25 positions x upright/inverted) and blank periods (10 trials), which were randomly ordered for each run and each participant. Each run began and ended with an additional 16s blank period. Throughout each run, participants performed a challenging RSVP letter detection task at the central fixation point (0.4° diameter). Participants’ fixation was monitored with an Eyelink 1000 eyetracker during the scan.

#### Trial structure

Each trial consisted of 7 faces (300ms on, 200ms off) and one blank interval (500ms) presented at a single spatial location. The same timing structure was used for the central letter presentation, including during no-face blank periods.

#### RSVP task

On each trial, participants were instructed to attend and respond to a stream of small, rapidly presented letters (A-S-D-F-G-H-J-K) and detect one-back repeats of the same letter. Half of all trials contained a repeat in the 7-letter stream, which occurred randomly in time within the trial. Letters were presented throughout the entire mapping run, and participants reported the task to be both difficult and attentionally engaging. Overall performance ranged from 77% to 98% correct (mean = 90%, SE = 2%), and we observed no difference between performance on trials that contained upright vs. inverted faces (*t*(11) = 0.531, p = 0.606), confirming that participants’ attention was not systematically drawn to either upright or inverted faces.

#### Face stimuli

We used front-view, grey-scaled face photographs previously used in Kay et al.^16^ and other work in the lab (VPNL-101-faces). These depicted 95 different individuals of various genders and races in a neutral expression; demographics represented that of the Stanford University undergraduate student body. Faces were presented in a circular aperture and were matched on the overall height of the head, resulting in substantial consistency in the position of internal facial features and some variability in the position and presence of external features, including hair.

### ROI definition

Each participant also completed an fMRI session of retinotopic mapping (4 runs) using brightly colored, 8Hz cartoon frames in a traveling wave aperture^18^, as well as 3-10 runs of a visual category localizer^57^. The localizer data was used to define face-selective regions in each participant by contrasting responses to images of child faces + adult faces versus images from eight other categories including bodies, limbs, objects, places, words and numbers, with a threshold of *t* ≥ 2.7, voxel level. Retinotopic regions V1-hV4 were defined from the retinotopy scans; boundaries between retinotopic areas were delineated by hand by identifying reversals in the phase of the polar angle map measurements and anatomical landmarks, as detailed in Witthoft et al.^58^. Face-selective region IOG was defined to exclude any voxels overlapping hV4.

### Data preprocessing

fMRI data was preprocessed following a standard pipeline using FSL (https://fsl.fmrib.ox.ac.uk/fsl/fslwiki/Fslutils) and mrVista (https://github.com/vistalab) tools. Data from the experimental runs and the ROI masks derived from the retinotopy and localizer analyses were co-registered to the participant’s high-resolution T1. Functional data underwent motion correction (within- and between-scans), slice-time correction (pRF mapping runs only), and low-pass filtering (60s). No spatial smoothing was done.

### pRF model fitting

We fit population receptive field (pRF) model estimates following the event-related paradigm introduced in Kay et al.^16^. This differs from many pRF implementations that fit the time course of responses to sweeping bars or rings/wedges (e.g. ^18,53^, and instead allows us to combine stimuli from different conditions (upright/inverted faces) in the same mapping run. mrVista functions (https://github.com/vistalab/vistasoft) were used to estimate GLM betas for each voxel in each of 50 conditions (25 locations x upright/inverted) using the SPM (http://www.fil.ion.ucl.ac.uk/spm) difference-of-gammas hemodynamic response function (HRF). Subsequent pRF modeling was done using these beta amplitudes.

Population receptive field estimates were made independently for each voxel in each mapping condition (upright/inverted) using the compressive spatial summation (CSS) model^29^ that assumes a circular 2D Gaussian pRF. The CSS model introduces a compressive power-law non-linearity (exponent *n* < 1) that increases position invariance within the pRF and better models responses in high-level visual areas^29^ including face-selective regions^16^. We estimated pRFs independently in each voxel and for each mapping condition by fitting six model parameters: gain, X, Y, σ, n. pRF size is defined as σ/√n, and visualized in coverage density maps with a contour at radius = 2 x size from the pRF center. Model fitting was performed using nonlinear optimization (MATLAB Optimization Toolbox).

For quantification of pRF properties, we selected voxels with a goodness-of-fit R^2^ > 0.2 in both the upright and inverted conditions; in pooled-voxel visualizations (**Figure 3**), a threshold of R^2^ > 0.5 was used for better visual clarity. Additionally, we trimmed voxels whose pRF fits were outside the range of the mapping stimulus (central 11°) or within the fixation (central 0.4°), as well as those whose size was estimated to be < 0.1°, as this typically corresponded to voxels that did not respond well to our stimuli (e.g. uniformly low responses across the visual field).

In the main experiment, face position was coded using a binary mask determined by the silhouette outline of the faces. These masks were drawn by hand for each face stimulus, and the model fit was based on an average of the specific faces that the individual participant saw at each of the 25 positions during his or her experimental session.

### Rescale + noise simulation (pRF experiment)

To elucidate how differential signal strength or model goodness-of-fit may have contributed to the observed differences in inverted- and upright-mapped pRF size and position estimates, we ran an iterative simulation on data from right hemisphere mFus-faces, pooled across participants. As shown in **Figure S2**, simulated data for each voxel was generated by taking beta estimates from the upright condition and matching beta range (‘rescaling’ step) and model goodness-of-fit (‘noise’ step) to that of the inverted condition. The rescaling step involved divisively scaling the range of the upright betas to match that of the inverted betas. The noise step iteratively added Gaussian noise to the betas until the simulated goodness-of-fit R^2^ was within .01 of the R^2^ in the inverted condition. To summarize the results of this simulation and compare between upright, inverted, and simulated-data estimates, we used bootstrapping (1000 draws) to estimate the median and 68% confidence interval of each model parameter of interest (Position Y, size, and gain).

### Quantifying pRF size vs eccentricity

pRF size linearly increases with eccentricity throughout the visual hierarchy^15–16,59^. Thus, we compared the size vs. eccentricity relationship for upright and inverted faces in each ROI. We performed this analysis in two ways: (i) pooling well-fitted voxels (R^2^ > 0.5) across participants in each ROI (**Figure 3**), and (ii) separately estimating the relationship for each participant and ROI (**Figure S3**). The former produces more robust line fits, while the latter provides an estimate of between-participant variability. Fitting was done by minimizing the L1 norm of the residuals, e.g., the sum of the absolute values of the residuals. This solution was chosen for its robustness to outliers in the underlying data.

### Visual field coverage

Visual field coverage density plots for each visual region in **Figures 5 and S4** were generated first for each participant, and then averaged across participants. Using a custom Matlab bootstrapping procedure, these density plots represent the proportion of pRFs in a region that overlap with each point in the visual field. Overlap is determined using a binary circular pRF at the estimated center, and with a radius of 2σ/√n^29^. This metric does not account for the Gaussian profile of individual pRFs, but allows for greater interpretability when combining data across pRFs and participants.

For each participant and ROI, density plots were generated by taking 1000 bootstrap samples of 80% of voxels with replacement. Each voxel was represented with a circular binary mask as described above, and coverage was computed by calculating the mean density across voxels for each bootstrap draw. The average of these bootstrapped images is taken as the density coverage for each participant. Participant-wise metrics, like FWHM area and center-of-mass (**Figure 5b**), are computed from these images. Averages across participant-wise images are taken as overall coverage density summaries (**Figure 5, Figure S6**); no rescaling or normalization is done, such that plotting colors retain meaningful quantitative information about pRF coverage across the visual field.

### Maximal-overlap location simulation for behavioral experiment

To evaluate the hypothesis that inverted-face recognition would be improved at the retinotopic location that produced maximal overlap with inverted-mapped pRFs, we ran a simulation using preliminary data from the first six participants in the main experiment (**Figure 6b, Figure S7**). We pooled voxels from bilateral mFus-faces across participants, and then took 1000 random samples of 100 voxels to generate a simulated pRF coverage. We then took the dot product of this coverage with an averaged absolute-contrast image of the internal features of the mapping faces at 3.2° diameter (such that the full face image, not the internal features, was 3.2°), positioned at 0.1° intervals spanning the central 3° × 3° of the visual field. For each simulated coverage map, this yielded an overlap metric at each of these 0.1° intervals (**Figure 6b, right panel**), and the maximal value was chosen. This yielded an average maximal overlap when the face was positioned at 0.67° to the left and 0.79° below the center; the leftward shift is a consequence of the fact that the right mFus-faces is larger than the left mFus-faces in most participants, so that there are more voxels that have pRFs covering the left than right visual field (see also^33^). For comparison, the same simulation using the internal features of an upright face and the upright-mapped pRFs yielded a maximal-overlap position at 0.48° to the left and 0.58° above the center. See also **Figure S7**, in which the maximal-overlap location is computed for each of the 12 subjects in the full experiment.

### Behavioral experiment

Following the simulation described above, we sought to probe recognition performance for upright and inverted faces at three positions: the *lower left* location determined to yield maximal overlap between inverted face features and inverted-mapped mFus-faces pRFs, a mirrored *upper right* location that is equivalently far from the center, but predicted to have much lower pRF overlap, and the *center* of the screen, where FIE is typically measured. The experiment followed a standard behavioral face inversion paradigm^23^, using a challenging recognition memory task. In brief, during the behavioral task (**Figure 7a),** participants fixated on a small central bullseye and saw a sequence of 3 faces each shown for 400ms, all upright or all inverted within each trial. After an 800ms interval, a target face appeared at the same location. Participants were asked to indicate whether the face was one of the faces in the prior triad (50% of trials) or not (50% of trials). The experiments were run in the eye tracking lab, and participants’ fixation was monitored with an Eyelink 1000 as described below. All 12 of the scan participants took part in this experiment after their scan session, and eye position was carefully monitored to ensure fixation. Data from 3 participants was excluded from the final analysis because they failed to maintain fixation on more than 25% of all trials. Performance was measured for each condition (location x face orientation) as a sensitivity index (d’).

#### Experimental design

Prior to starting the behavioral experiment, participants received practice on the task with performance feedback. Following the practice period, each participant completed 18 blocks of 20 trials. Position varied across blocks, such that each position (lower left, center, upper right) was probed on 6 blocks. Within each block, half of the trials used upright faces and half used inverted faces; trial ordering (i.e., inversion) was randomized. In total, each condition (face orientation x position) was probed in 60 trials, and the experiment took ~30 minutes to complete.

#### Trial structure

Each trial (**Figure 7**) consisted of 3 faces presented for 400ms, followed by an interstimulus interval of 800ms, and a 400ms target face. Participants were asked to respond via keyboard whether the target face, which was marked with a thin green outline, had appeared in the previous triad. Participants were given 2400ms from the appearance of the target face to respond. Targets appeared on half of trials, randomly as the first, second, or third face in the triad. On trials when the target was absent, it was replaced by a distractor face that was matched in general appearance (e.g., ‘blonde female with long hair’) to the target, as maximally allowed by our set of 95 face identities. Face identities were the same as in the scan experiment. In the behavioral experiment, we used front-view faces and also ± 15° viewpoint angle images of the same individuals to increase the difficulty of the task. Trials were controlled so that the target image and its incidence in the triad were always of different viewpoints. That is, an identical image was never used in the triad and as the target. Each trial was preceded by a 2s fixation intertrial interval (ITI) during which eye position was recorded to be used toward drift correction.

#### Eyetracking

Eyetracking during the behavioral experiment was performed using an Eyelink 1000 with a sampling rate of 1000Hz, and analyzed using a custom Matlab pipeline. Prior to starting the experiment, participants received practice on the task with both performance feedback and realtime eyetracking feedback: we plotted their current eye position as a series of red dots while they performed the practice task at each of the probe positions. Participants were required to achieve good fixation (< 0.5° deviation across trials in the second half of the practice) before they could proceed to the main task. Additionally, to improve the validity of our eyetracking metrics, we performed drift correction during each ITI; participants were aware of this and were instructed to keep steady fixation even between trials. This allowed us greater confidence in the absolute values of the tracked eye position on our displays, which was contained in the central ~5° of the visual field in total. Following the experiment, we removed from the analysis any trials on which participants broke fixation, and excluded 3 participants for whom this happened on > 25% of all trials. For the remaining 9 participants, an average of 9.51% (s.d. = 4.65%) of trials across conditions were removed as a result of breaking fixation.

## Acknowledgements

This work was supported by NSF BCS #1756035 to KGS and SP.

## Author Contributions

SP, KK, and KGS designed the experiment. SP and DF collected and analyzed data. SP and KK wrote the analysis pipeline. KGS oversaw the data analyses. SP, KK, DF, and KGS wrote the manuscript.

## Data and Code Availability

Upon publication, data and code will be publicly available at github.com/VPNL/invPRF.

## Supplementary Information

**Figure S1.**
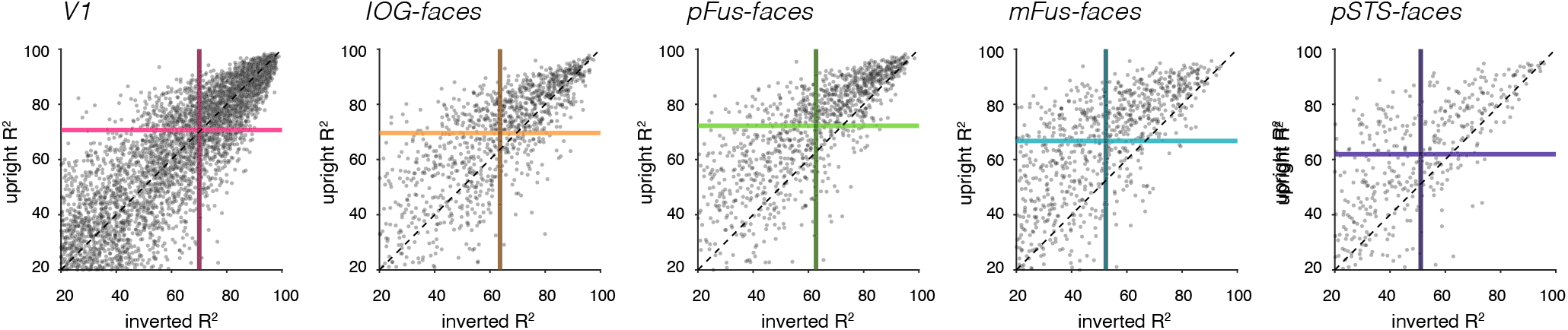
Voxelwise comparison of model goodness-of-fit (R^2^) across the visual hierarchy. Voxels are pooled across subjects, and the model goodness-of-fit for each voxel in the data set is shown. Colored lines indicate the mean R^2^ across voxels in the upright (horizontal, light colors) and inverted (vertical, dark colors) mapping conditions. Model goodness-of-fit was equivalent across mapping conditions in V1, but was lower in the inverted face condition in all face-selective areas (see also **Table S1**).

**Figure S2.**
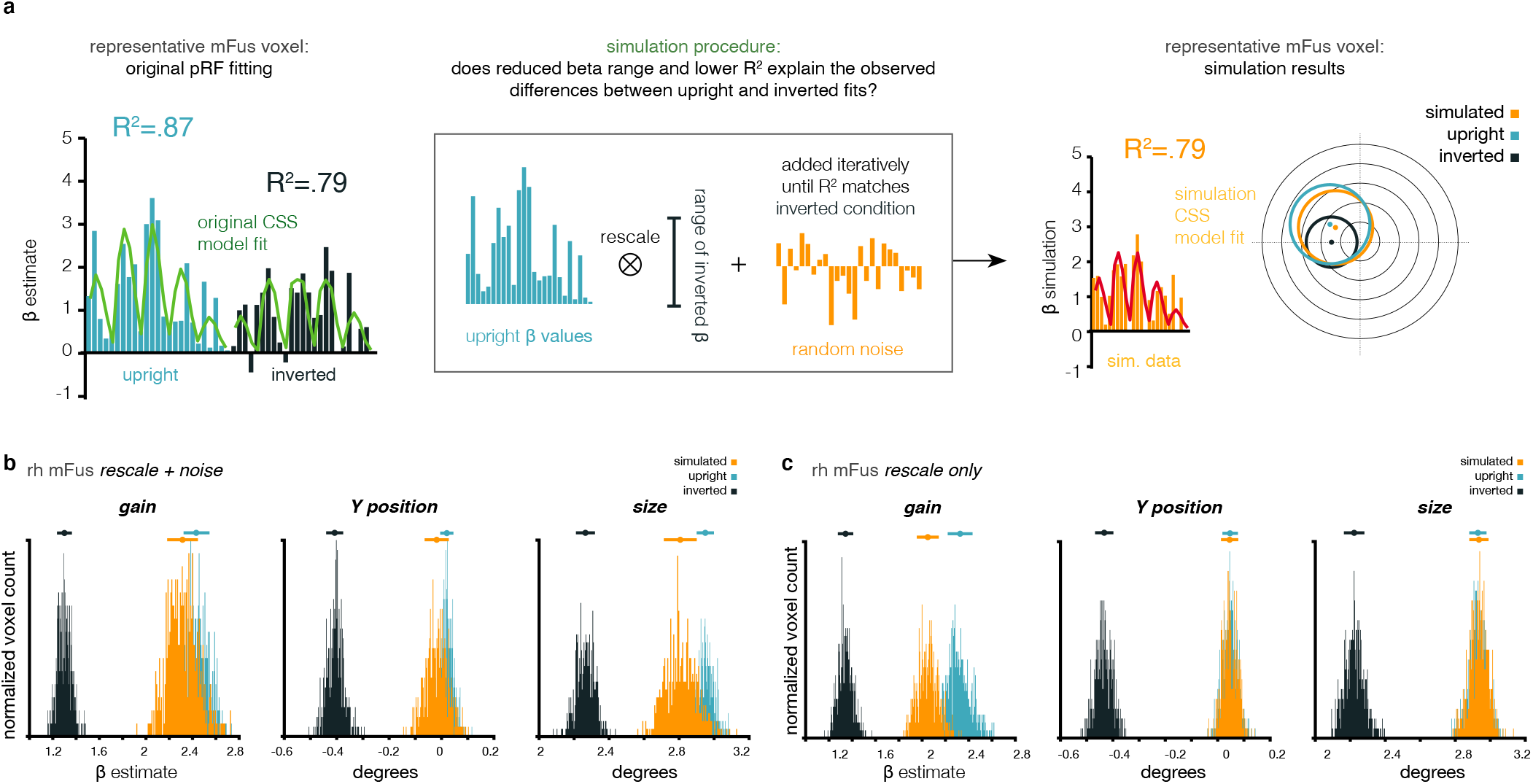
Iterative noise simulation procedure and results. **(a)** To what degree do the observed differences in signal strength and R^2^ between inverted and upright faces contribute to differences in pRF position and size estimates? To determine this, we ran a simulation over mFus-faces voxels: for each voxel, we took the 25 betas from the upright mapping condition (light blue), rescaled them to match that the range of the 25 betas from the inverted mapping condition (dark blue; ‘rescale’ step), and then iteratively added random noise (‘noise’ step) until the simulated model fit R^2^ was equal to the model R^2^ in the inverted condition. In this example voxel, the simulated pRF fit (orange) much more closely resembles the upright than the inverted condition fits. **(b)** Bootstrapped parameter estimates for the rescaling + noise procedure described in **(a)**. While the simulated fits do show some differences in size and gain compared to upright pRFs, the properties of the estimated pRFs much more resembled the upright than inverted pRF and differences between upright and simulated pRFs are substantially smaller in magnitude than what we observe for inverted faces, despite matching R^2^ and signal level (*βs*). *Horizontal line markers* above distributions indicate median and 68% confidence intervals around the estimates. **(c)** Results of an additional simulation which was the same as (b) except that we omitted the noise step. Changing the response amplitude has minimal effect on the size or position of pRFs. Together, these simulations allow us to conclude that differences in model goodness-of-fit and weaker signal in response to inverted faces cannot fully account for the observed differential pRF estimates for upright and inverted faces in the main experiment.

**Figure S3.**
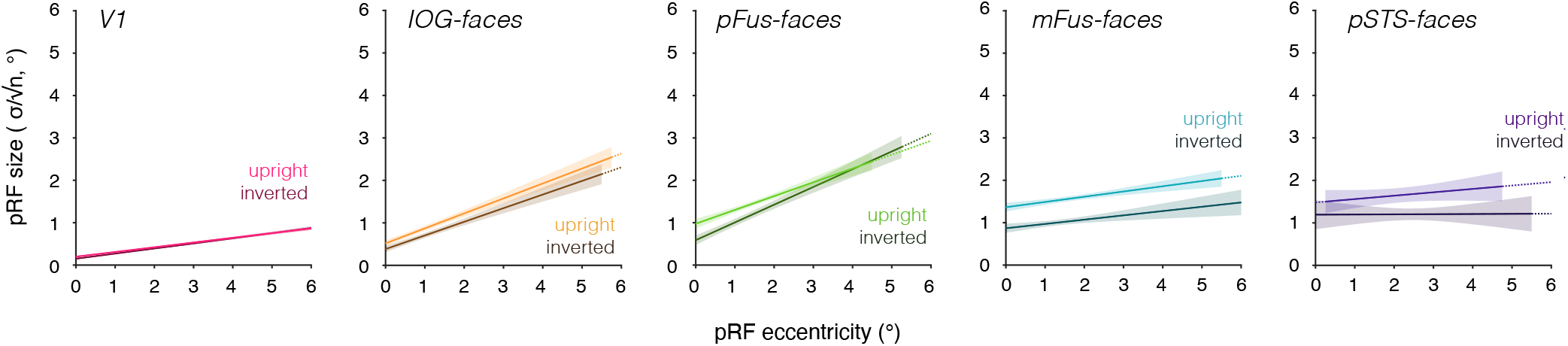
Inverted-mapped pRFs are smaller in face-selective regions across eccentricity. Related to **Figure 3**, here the size by eccentricity relationship in each ROI is estimated for each individual participant to provide an estimate of between-participant variability. Lines are generated by fitting least-squares lines to individual participant data and then averaging these fits. The line is subsampled at 0.25°, and shaded regions indicate ± SEM of the estimated model across participants, at eccentricities containing > 0 voxels in each individual participant. Decreased size for inverted-mapped pRFs is most apparent in mFus-faces, but is also evident in pFus- and pSTS-faces. We observed no reliable size differences across participants in V1 or IOG-faces. See also **Table S2**.

**Figure S4.**
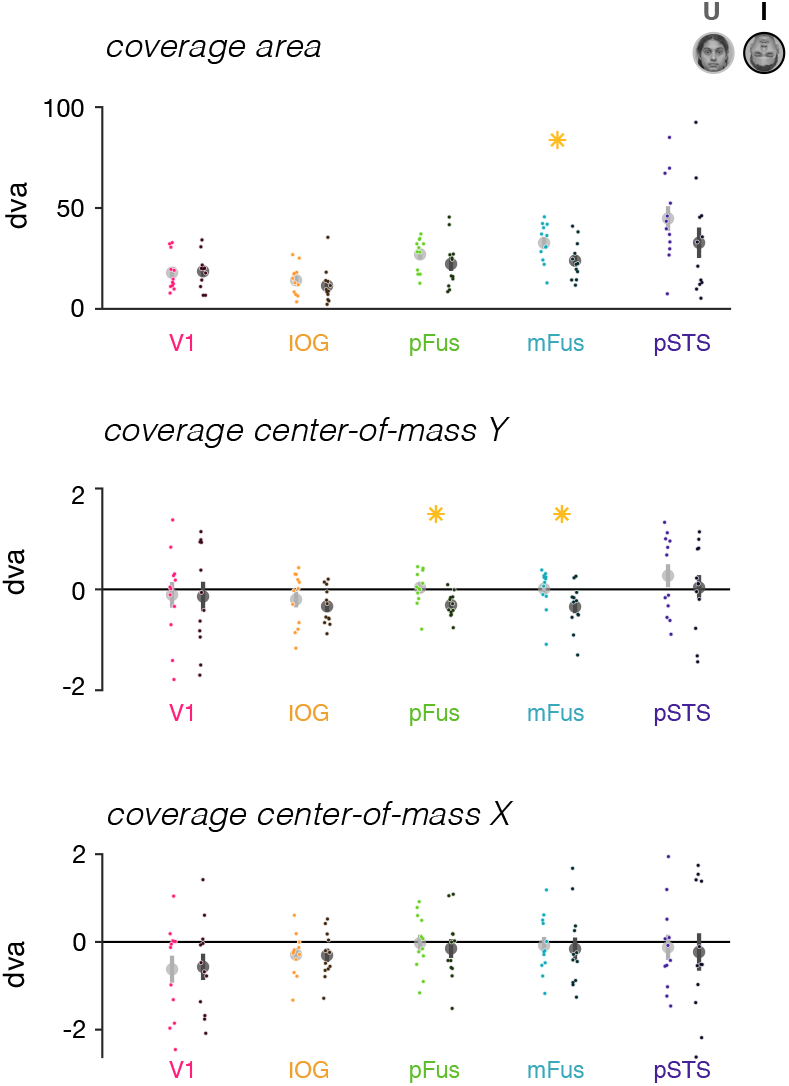
Coverage metrics across participants. Related to **Figure 5**; we estimated for each ROI and participant the area of coverage and the center-of-mass of the coverage. Area (top) was computed for the region at full-width-half-maximum of the coverage plots shown in **Figure 5**. The center of mass is taken over the same region, and has two parameters: a vertical Y coordinate (middle) and horizontal X position (bottom). *Dots*: individual participant metrics. *Asterisks*: significant differences between upright and inverted face conditions at p < .05, paired t-test.

**Figure S5.**
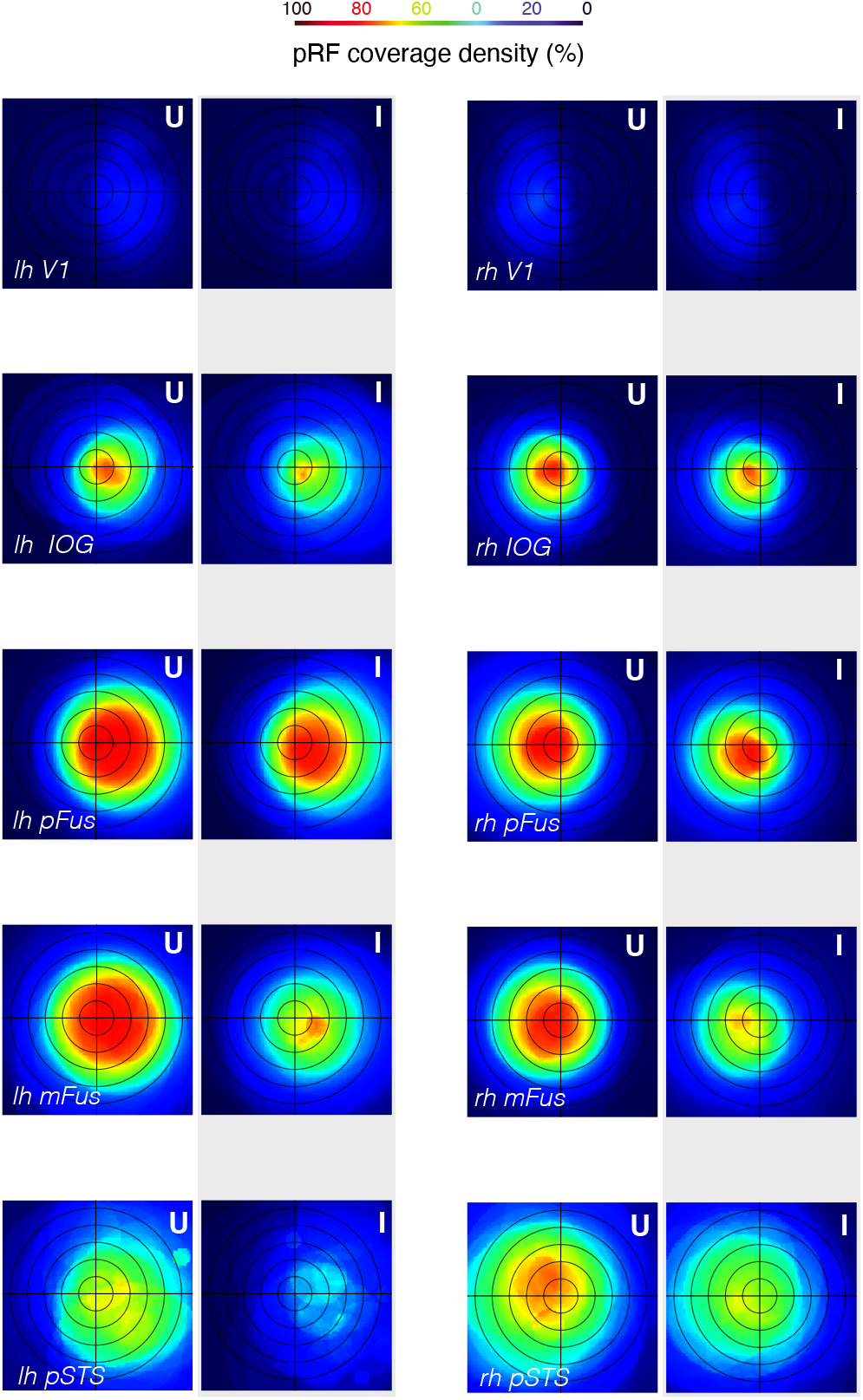
Average pRF coverage density across participants by hemisphere. Plotting conventions follow **Figure 5**; coverage density approximates the proportion of pRFs that overlap a given region in space. Mapping with upright faces (U) replicates previously reported properties of coverage in face-selective regions, including a strong contralateral representation of the fovea in pFus- and mFus-faces^1,2^, and less foveal bias in pSTS-faces than in the ventral regions^3^. Mapping with inverted faces (I) yields significantly smaller full-width-half-max coverage area in lh-mFus, rh-mFus, lh-pSTS (*t*’s(11) > 2.30, p’s < 0.044) and a significant downward shift of the center-of-mass of the coverage in lh-pFus, rh-pFus, rh-IOG (*t*’s(11) > 2.33, p’s < 0.041). There were no significant differences in the horizontal location of the center-of-mass in any region and no significant changes to any of these parameters in either lh- or rh-V1.

**Figure S6.**
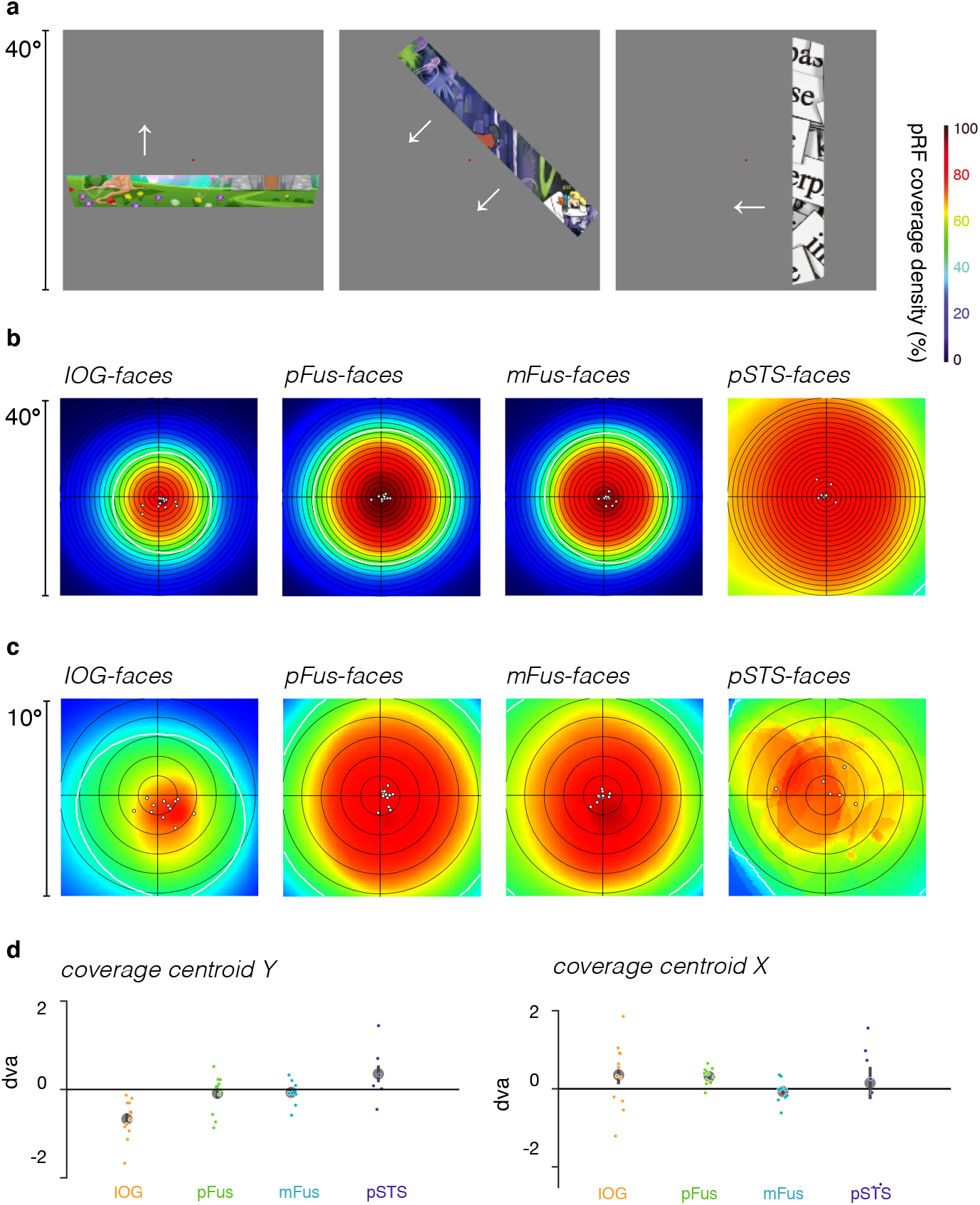
pRF mapping with cartoon stimuli yields pRF coverage consistent with upright-face mapping. 12 participants in the current study took part in a separate pRF mapping experiment that used completely different mapping stimuli, mapping timing, and fitting procedures ^3^, allowing us to evaluate whether any non-optimal stimulus would yield a similar reported pattern of results as inverted faces, or whether changes in pRF properties and coverage are specific to face inversion. For brevity, here we refer to ‘toonotopy data,’ as the subset of data from Finzi et al.^3^ for the participants in the current study (N = 12), analyzed using the same participant-wise ROIs as in the current study. Unlike the current study, for the toonotopy data we fit the CSS pRF model over mean BOLD timecourses (procedure: http://cvnlab.net/analyzePRF) rather than to estimated beta weights for discrete mapping stimuli. To better compare the widefield data to the current study, we analyzed both the full toonotopy datasets of the 12 subjects, and also a subset of the data matched to the spatial extent of the mapping stimuli in the current study (eccentricity < 5°, size < 10°). **(a)** Toonotopy mapping stimuli were comprised of rapidly-presented cartoon frames (8Hz presentation) within a bar aperture that swept the visual field in 8 directions, and covered the central 40° of the visual field over the course of an fMRI run. **(b)** For the 12 participants in the current study, the average pRF coverage in face-selective ROIs from the full toonotopy data. As in Figure 5, *white contour*: the average FWHM of coverage density. *White dots*: individual participants’ FWHM center-of-mass position. As expected, the widefield nature of the toonotopy stimuli of Finzi and colleagues ^3^ allow us to measure pRFs beyond the ~10° extent of the mapping stimuli used in our main experiment. To draw better comparison to the current study, **(c)** presents equivalent coverage plotting of the subset of subjects’ pRFs in the toonotopy data matched to the ~10° extent of the mapping stimuli of the current study. Cross-participant metrics of this coverage in the face-selective regions are shown in **(d)**. While we cannot quantitatively compare visual field coverage area between the two studies due to the differing spatial extent of the mapping stimuli, we see qualitatively similar coverage in response to the toonotopy mapping stimuli and the upright face mapping in the current study. That is, the visual field coverage was foveally biased in ventral face-selective regions and progressively increased from IOG-to pFus-to mFus-faces. We see no evidence of a downward shift in the center-of-mass for pFus- and mFus-faces as observed with inversion.

**Figure S7.**
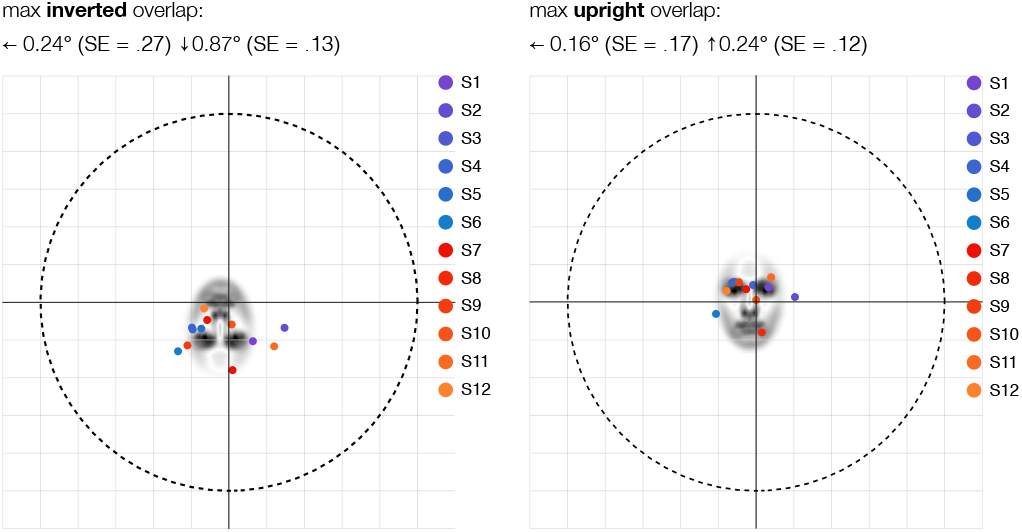
Positions in the visual field yielding maximal pRF overlap for each of our participants. As described in **Figure 6**, we sought to determine the position in space at which the internal features of an 3.2° upright face (left panel) maximally overlapped with upright-mapped pRFs, and the position in space at which the internal features of an inverted face (right panel) maximally overlapped with inverted-mapped pRFs. The bootstrapping simulation used to determine the positions at which faces were shown in the behavioral experiment was based on pooled preliminary data from the first 6 participants in the experiment (*S1-6*, cool colors). We repeated the simulation procedure on an individual-participant basis (1000 draws per participant, 80% of voxels sampled with replacement), including the participants whose pRF data was collected after the initial simulation (S7-12, warm colors). *Dots*: represent bootstrapped estimates for individual participants; *face images* in each panel are shown at the mean maximal overlap position in each condition. Overall, while some inter-participant variability is present, the mean positions determined across all participants align well with the position used in the behavioral experiment, such that maximal overlap for inverted faces occurs in the lower left visual field, while maximal overlap for upright faces occurs is more central and trends slightly to the upper left.

**Figure S8.**
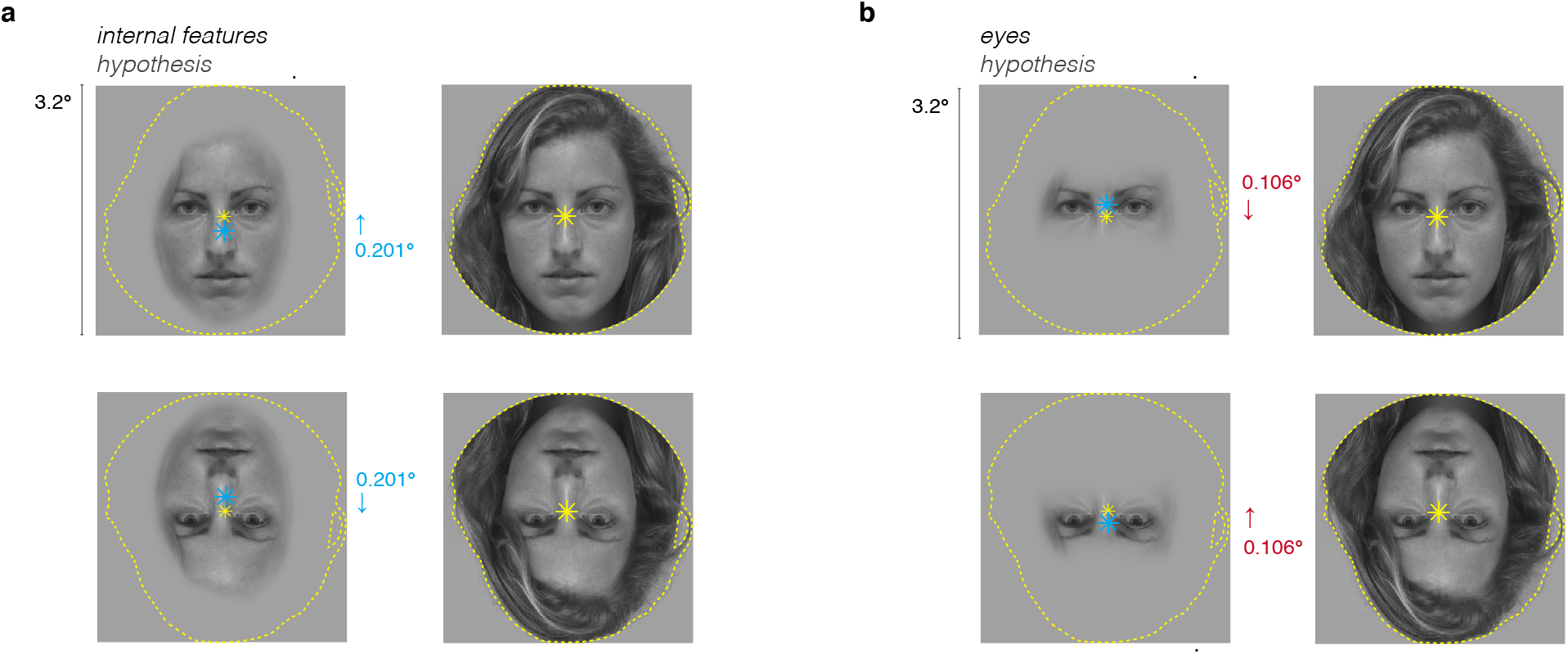
The location of internal face features may predict the magnitude and directionality of the reported pRF shifts with face inversion. In this schematic, we present two hypotheses that examine how the location of internal features (left) or eyes alone (right) may contribute to the positional shifts in pRFs due to face inversion. In the pRF model, each individual face location is coded as a binary silhouette of the full-face image that participants saw, here shown via *yellow dashed line*. Because pRF model fitting relies on computing the dot product of a hypothesized Gaussian pRF and this binary image, we can infer that the center-of-mass of the binary silhouette is an important factor in determining the estimated pRF position. The weighted centroid of the binary silhouette for the sample image is marked by *yellow asterisks* in all panels. To account for the observed shift in Y position in response to face inversion, we evaluated the **(a)** *Internal features hypothesis*: the position of the internal face features, rather than the full face image, drives spatial responses in face-selective regions. This can be approximated by calculating the weighted centroid of the internal features of this face for the upright and inverted conditions (left panel, *blue asterisks*). Correspondingly, if the internal features rather than the entire face determine the location of the pRF in both the upright and inverted mapping conditions (blue), our implemented pRF model (yellow) will yield position estimates ~0.2° above the location of the internal features of the face in the upright condition, and ~0.2° below in the inverted condition. The overall consequence of these offsets would be a ~0.4° lower pRF estimate in the inverted condition than in the upright condition, which is similar to the magnitude and directionality of effects we observed in pFus- and mFus-faces. **(b)** *Eyes hypothesis*: the position of the eyes alone, rather than the full face image, drives spatial responses in face-selective regions. This can be approximated by calculating the weighted centroid of the eyes of this face for the upright and inverted conditions (right panel, *blue asterisks*). Correspondingly, if the location of the eyes within the mapping stimulus drives spatial responses in a voxel, we would expect to see (i) smaller difference between the upright- and inverted-mapped pRF estimates, as the centroid of the eye region of our face images (blue asterisk) is typically well-aligned with the centroid of the binary silhouette (yellow), and (ii) inverted-mapped pRFs that are shifted upward ~0.2° relative to the upright-mapped condition. This is a poorer match to the effects of face inversion observed experimentally; neither manipulation predicts the observed differences in pRF gain or size.

**Table S1.** Results of three-way repeated measures analyses of variance (ANOVA) with factors of hemisphere (right/left), region of interest (IOG-/pFus-/mFus-/pSTS-faces), and condition (upright/inverted) for each pRF mean parameter estimate across participants. One participant had insufficient lh-pSTS data (0 voxels above threshold) and was excluded from this analysis, leaving N = 11. Related to **Figure 2.**

| center position Y estimate |  |  |
| --- | --- | --- |
| significant main effects: |  |  |
| condition | F(1,10) = 8.82 | p = 0.014 |
| significant interactions: |  |  |
| none |  |  |
| not significant: |  |  |
| hem | F(1,10) = 0.10 | p = 0.758. |
| ROI | F(3,30) = 0.10 | p = 0.681 |
| hem x ROI | F(3,30) = 0.55 | p = 0.649 |
| hem x condition | F(1,10) = 2.61 | p = 0.137 |
| ROI x condition | F(3,30) = 0.83 | p = 0.489 |
| hem x ROI x condition | F(3,30) = 0.67 | p = 0.577 |

| center position X estimate |  |  |
| --- | --- | --- |
| significant main effects: |  |  |
| hem | F(1,10) = 375.60 | p < 0.001 |
| significant interactions: |  |  |
| hem x ROI | F(3,30) = 4.49 | p = 0.010. |
| not significant: |  |  |
| ROI | F(3,30) = 2.15 | p = 0.114 |
| condition | F(1,10) = 0.31 | p = 0.592 |
| hem x condition | F(1,10) = 1.16 | p = 0.306 |
| ROI x condition | F(3,30) = 0.39 | p = 0.763 |
| hem x ROI x condition | F(3,30) = 0.06 | p = 0.982 |

| size estimate ($\sigma / \sqrt{n}$) | | |
| --- | --- | --- |
| significant main effects: |  |  |
| ROI | F(3,30) = 7.13 | p = 0.001 |
| condition | F(1,10) = 56.07 | p < 0.001 |
| significant interactions: |  |  |
| hem x ROI | F(3,30) = 3.36 | p = 0.032 |
| ROI x condition | F(3,30) = 7.53 | p = 0.001 |
| not significant: |  |  |
| hem | F(1,10) = 0.01 | p = 0.920. |
| hem x condition | F(1,10) = 1.18 | p = 0.303 |
| hem x ROI x condition | F(3,30) = 2.64 | p = 0.068. |

| gain estimate |  |  |
| --- | --- | --- |
| significant main effects: |  |  |
| none |  |  |
| significant interactions: |  |  |
| ROI x condition | F(3,30) = 4.66 | p = 0.009 |
| not significant: |  |  |
| hem | F(1,10) = 2.77 | p = 0.127 |
| ROI | F(3,30) = 0.61 | p = 0.615 |
| condition | F(1,10) = 2.07, | p = 0.181 |
| hem x ROI | F(3,30) = 0.75 | p = 0.530 |
| hem x condition | F(1,10) = 0.05 | p = 0.824 |
| hem x ROI x condition | F(3,30) = 0.36 | p = 0.781 |

| model R<sup>2</sup> |  |  |
| --- | --- | --- |
| significant main effects: |  |  |
| ROI | F(3,30) = 17.59 | p < 0.001 |
| condition | F(1,10) = 80.36 | p < 0.001 |
| significant interactions: |  |  |
| ROI x condition | F(3,30) = 7.47 | p = 0.001 |
| not significant: |  |  |
| hem | F(1,10) = 0.19 | p = 0.669 |
| hem x ROI | F(3,30) = 1.13 | p = 0.352 |
| hem x condition | F(1,10) = 1.28 | p = 0.283. |
| hem x ROI x condition | F(3,30) = 0.23 | p = 0.872 |

**Table S2.** Results of two-way repeated measures ANOVA with factors of ROI (IOG-/pFus-/mFus-/pSTS-faces) and condition (upright/inverted) on the relationship of size and eccentricity in bilateral face-selective areas (related to **Figure S3**) across participants (N=12).

| fit line slope |  |  |
| --- | --- | --- |
| significant main effects: |  |  |
| ROI | $F(3) = 3.23$ | $p = 0.035$ |
| significant interactions: |  |  |
| *none* |  |  |
| not significant: |  |  |
| condition | $F(1) = 0.04$ | $p = 0.853$ |
| ROI x condition | $F(3) = 0.45$ | $p = 0.719$ |

| fit line intercept |  |  |
| --- | --- | --- |
| significant main effects: |  |  |
| condition | $F(1) = 9.08$ | $p = 0.012$ |
| significant interactions: |  |  |
| *none* |  |  |
| not significant: |  |  |
| ROI | $F(3) = 2.80$ | $p = 0.055$ |
| ROI x condition | $F(3) = 0.58$ | $p = 0.633$ |

**Table S3.** Results of two-way repeated measures ANOVA on the individual participants’ (N=12) coverage density metrics in bilateral face-selective areas (related to **Figure 5**).

| coverage density FWHM area |  |  |
| --- | --- | --- |
| significant main effects: |  |  |
| ROI | F(3) = 3.10 | p = 0.040 |
| condition | F(1) = 7.21 | p = 0.021 |
| significant interactions: |  |  |
| none |  |  |
| not significant: |  |  |
| ROI x condition | F(3) = 0.39 | p = 0.758 |

| center of mass Y position |  |  |
| --- | --- | --- |
| significant main effects: |  |  |
| ROI | F(3) = 3.10 | p = 0.040 |
| condition | F(1) = 7.21 | p = 0.021 |
| significant interactions: |  |  |
| none |  |  |
| not significant: |  |  |
| ROI x condition | F(3) = 0.39 | p = 0.758 |

| center of mass X position |  |  |
| --- | --- | --- |
| significant main effects: |  |  |
| none |  |  |
| significant interactions: |  |  |
| none |  |  |
| not significant: |  |  |
| ROI | F(3) = 0.18 | p = 0.907 |
| condition | F(1) = 0.25 | p = 0.627 |
| ROI x condition | F(3) = 0.05 | p = 0.984 |

